# Direct Optical Detection of Factor Xa Activity in Minimally Processed Whole Blood

**DOI:** 10.1101/2025.02.07.636760

**Authors:** Alyssa P. Cartwright, Benjamin C. Wollant, Elizabeth S. York, Liwei Zheng, Steven Yee, Huong C. Chau, Glaivy Batsuli, H. Tom Soh

**Author notes:** Equal contribution (Alyssa P. Cartwright, Benjamin C. Wollant).

## Abstract

The ability to measure factor Xa activity directly in whole blood samples offers a path toward point-of-care monitoring and personalized anticoagulant dosage, potentially reducing bleeding risk and other anticoagulant-associated complications. We present a strategy to enable direct optical detection of factor Xa in minimally processed whole blood samples. Our strategy relies on a custom FRET-pair labeled DNA-peptide substrate, allowing FRET ratio to be monitored as an indicator of factor Xa activity. Substrates are tethered to a tapered-fiber sensor to allow evanescent detection of fluorescence directly at the sensor surface, minimizing background media interference and enabling detection directly in blood samples. After characterizing the custom substrate and demonstrating the correlation of fiber-based measurements to an existing chromogenic assay, we demonstrate the detection of endogenous factor Xa activity in >85% whole blood. Finally, we demonstrate the detection of therapeutic concentrations of enoxaparin, a widely used anticoagulant, directly in 90% whole blood in less than an hour and correlate these measurements to activated partial thromboplastin time (aPTT) testing. Together, these results indicate a promising strategy to achieve point-of-care factor Xa detection, enabling personalized anticoagulant treatment and reducing adverse outcomes.

## Introduction

Anticoagulant therapy is widely used in the treatment and management of numerous conditions related to dysregulated hemostasis, including myocardial infarction, stroke, and venous thromboembolism [1]. In 2019, over $7.38B was spent on oral anticoagulants under Medicare prescription drug coverage for 5.2M beneficiaries [2] with total spending and usage rising since [3]. Many conditions treated with anticoagulants are linked to the dysregulation or dysfunction of specific clotting factors within the coagulation cascade, which in turn can lead to thrombosis [4]. Factor Xa is a key coagulation enzyme essential for thrombin generation, linking the upstream events of the intrinsic and extrinsic pathways to the final steps of fibrin clot formation [4, 5]. Factor Xa thus serves as a useful target for antithrombotic drugs, as its inhibition prevents coagulation triggered from either pathway [6]. Antithrombotic agents targeting factor Xa include direct oral anticoagulants (DOACs) that specifically target factor Xa as well as subcutaneously injected low molecular weight heparins (LMWH) that inhibit serine proteases including factor Xa [6, 7]. Despite the widespread use of anticoagulants, bleeding complications remain the primary risk of anticoagulant administration [8, 9], as described in the Joint Commission’s National Patient Safety Goals [10]. Dosing management within the therapeutic window to mitigate bleeding complications is important for anticoagulants, including those that target factor Xa. For commonly used DOACs, the availability of antidotes remains limited in most clinical settings [11]. Furthermore, enoxaparin, the most commonly used LMWH for venous thromboembolism in pediatric patients [12, 13, 14], is less effectively reversed by protamine, the standard antidote for unfractionated heparin [15]. Achieving accurate enoxaparin dosage is further complicated by variable drug pharmacokinetics in pediatric populations, necessitating frequent testing of anti-factor Xa activity [12]. Direct detection of factor Xa activity at the point-of-care may enable personalized dosage for enoxaparin in pediatric populations. Unfortunately, current methods for measuring anticoagulant effect face key limitations in molecular specificity and sample preparation requirements that hinder point-of-care usage for guidance of antithrombotic drug administration. Standard time-to-clot assays, such as activated partial thromboplastin time (aPTT) and prothrombin time (PT) assays, do not provide information on specific clotting factor activity and require processing blood into plasma, complicating efficiency and specificity for point-of-care usage [16, 17, 18]. Factor-specific versions of time-to-clot (one-stage) coagulation assays do exist but are typically less sensitive to common pathway factors, like factor Xa, that are common therapeutic targets [19, 20, 21]. Chromogenic assays, employing a color-changing substrate that is cleaved by a specific coagulation factor, can directly measure factor Xa activity, providing an indicator of the effect of factor Xa-inhibiting drugs like enoxaparin. However, these assays require blood processing and costly reagents, are time-consuming to run manually, and are difficult to automate [22, 23]. Thus, the technology to rapidly and accurately monitor factor Xa activity in point-of-care settings, providing specific molecular insights at the point of drug action, would aid clinicians in real-time drug monitoring and dosing, allowing safer, personalized dosing regimens of antithrombotic therapies [9, 24].

To address this need, we have combined a sensitive fiber-optic sensor with a fluorescent substrate to achieve real-time direct optical detection of factor Xa activity in small volumes (∼50 µL per replicate) of minimally processed whole blood. We employ a custom peptide-DNA substrate with a Förster resonance energy transfer (FRET)-pair of fluorophores conjugated on opposite sides of a known factor Xa cleavage site. Enzymatic cleavage of the substrate separates the FRET pair, resulting in a measurable decrease in FRET efficiency. Notably, the use of FRET-based, ratiometric measurement insulates our detection mechanism from matrix effects in whole blood that may modulate fluorescence and confound single fluorophore measurements. Our substrate is tethered to a fiber-optic evanescent sensor previously developed by our group, which selectively measures fluorescence signals occurring within a few hundred nanometers of the fiber surface [25]. Our system can resolve FRET signal changes resulting from specific factor Xa activity at the fiber surface even in the optically complex environment of whole blood while minimizing artifacts from scattering, absorbance, and autofluorescence in the bulk sample volume. After describing the design, synthesis, and validation of the custom FRET-based substrate in buffer, we demonstrate surface-based detection of known factor Xa concentrations in a high-protein-content artificial serum environment and correlate these measurements to results from a standard chromogenic assay. Finally, we use this system to measure endogenous factor Xa activity in >85% whole blood. As a sample application, we demonstrate that we can differentiate between normal, endogenous factor Xa activity and factor Xa activity that has been altered by treatment with therapeutic levels of enoxaparin within 60 minutes and validate these results via correlation to an aPTT assay. These results show that our sensor is capable of rapid measurement of factor Xa activity directly in minimally processed blood, providing specific molecular insight at the site of drug action as a promising tool for guiding and monitoring anticoagulant therapies at the point-of-care.

## Results and Discussion

### System overview

Our factor Xa substrate molecule (**Fig. 1a**) features a FRET pair of fluorophores separated by a polypeptide sequence containing a known factor Xa cleavage site, as well as a biotin tag to enable streptavidin-based tethering to the surface of our fiber-optic probe. To assemble our substrate, we began with a single-stranded DNA sequence comprising a 24-nucleotide (nt) poly-T sequence with a 5’ dibenzocyclooctyne (DBCO) moiety, an internal Cy3 label (the donor fluorophore) on the 6^th^ nucleotide, and a 3’ biotin group. The poly-T sequence is highly hydrophilic, improving the solubility of the substrate. We then performed a click chemistry reaction to couple the 5’ DBCO group to a 13-amino acid (aa) polypeptide sequence containing a C-terminal azide modification as well as a factor Xa cleavage site (IEGR) and N-terminal ATTO 643 label (the acceptor fluorophore; **SI, Note 1, Table S1, Fig S1**). The short length of the polypeptide sequence keeps the two fluorophores in sufficient proximity to allow efficient FRET.

**Figure 1.**
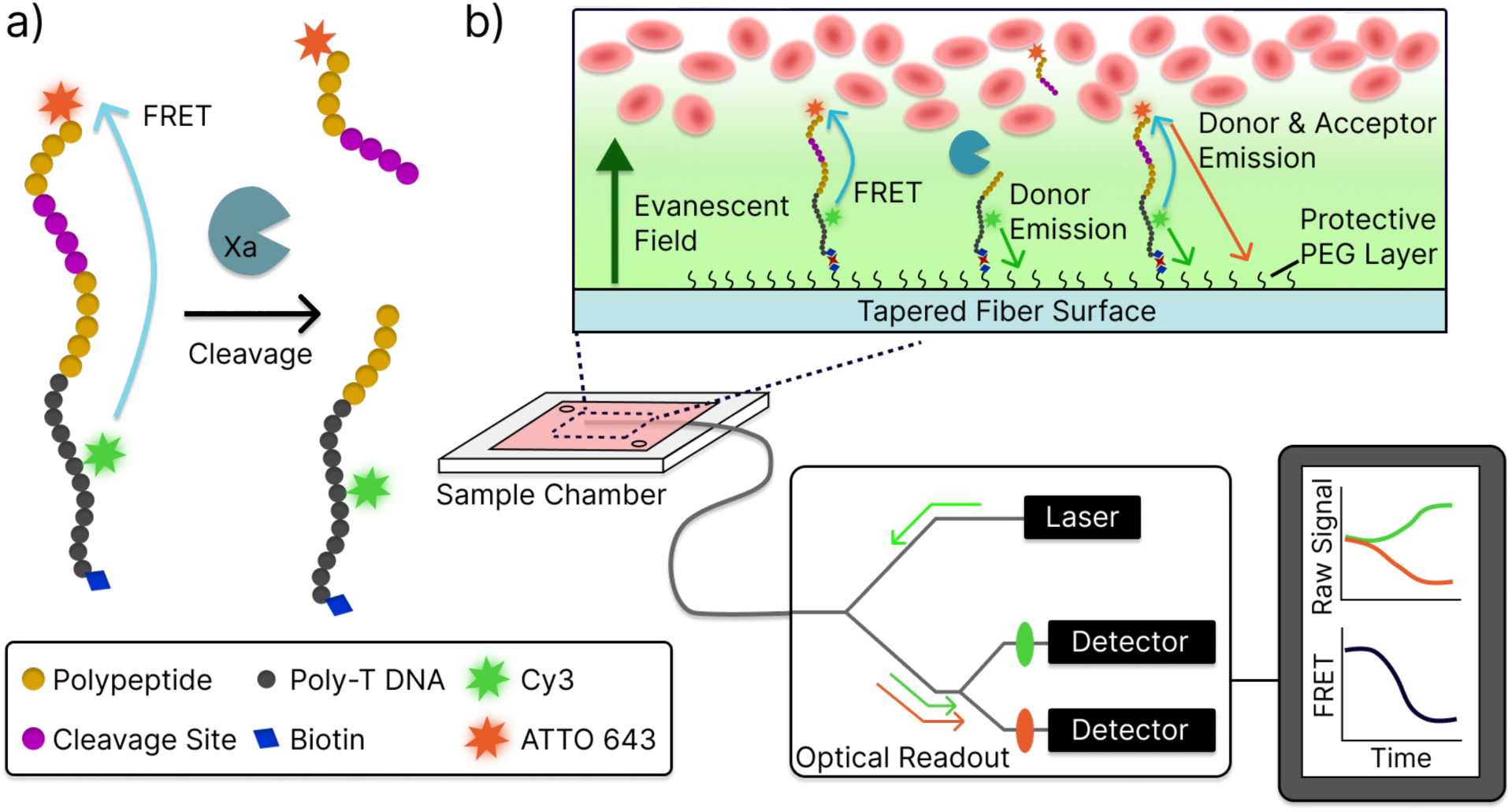
Design of our factor Xa activity sensor. **a**) A FRET-labeled substrate for detecting factor Xa activity. This substrate consists of a 23-nt poly-T DNA strand (black) with a terminal biotin group (blue diamond) and internal Cy3 label (green star), which has been coupled via click chemistry to a 13-aa polypeptide (gold) containing the factor Xa cleavage site (IEGR, magenta) and a distal ATTO 643 label (orange star). Figure not to scale. **b**) This substrate is tethered to the surface of a tapered optical fiber-based evanescent sensor, which can monitor changes in both donor and acceptor fluorescence to calculate FRET change. Because it relies on evanescent sensing, this system is robust against background interference from scattering, absorbance, and autofluorescence in blood samples. The majority of the fiber surface is coated with a protective PEG layer to reduce biofouling, and a small percentage of these PEG molecules contain a terminal biotin. These biotin groups are used to subsequently coat the surface with neutravidin, which is used to tether biotinylated substrates to the fiber probe. Figure not to scale.

We achieve sensitive detection in whole blood by combining evanescent detection with ratiometric FRET-based measurements. Specifically, we coupled our fluorescent substrate to a previously developed fiber-optic probe platform (**Fig. 1b**) [25], which employs a tapered optical fiber to detect fluorescent signals in close proximity to the fiber tip. When light from a laser source is coupled into this tapered fiber probe, an evanescent field is established at the tip, which decays exponentially in strength as the distance from the fiber surface increases. This means that the sensing region is confined to a few hundred nanometers radially from the surface. Within this region, surface-bound fluorophores are efficiently excited by the incident light, and a portion of the subsequently emitted fluorescence is coupled back into the fiber probe. Beyond this narrow region, the exponential decay of the evanescent field greatly reduces the efficiency of both excitation and detection of fluorescence in the bulk sample volume. This effect suppresses potential artifacts arising from scattering and absorbance due to red blood cells or competing signals from autofluorescent species in the blood [26], making it possible to achieve direct detection of factor Xa activity in whole blood. The collected fluorescence is transmitted via the optical fiber to a pair of single photon counting module (SPCM) detectors that enable sensitive measurement of the donor and acceptor signals. Additionally, the use of ratiometric FRET-based measurements further ensures that our sensor is robust against environmental effects that can interfere with single-color fluorescent assays. Fluorophore emission is sensitive to the local environment, including changes in temperature and pH, and can even be partially quenched by red blood cells [27]. These factors can confound single color measurements, as fluorescence changes caused by non-specific environmental factors cannot easily be distinguished from the true signal. By using a FRET pair, we effectively normalize these background-associated shifts by using the ratio of two fluorescent channels as the readout.

### Validating detection of factor Xa activity

We first set out to demonstrate that our substrate yielded a measurable change in FRET efficiency upon factor Xa cleavage and that this change was proportional to the concentration of factor Xa. With 13 aa and five nt separating the two fluorophores, we expected the FRET pair to be within a few nanometers in the fully assembled substrate, yielding a strong FRET signal (Förster radius ∼4.5 nm). Cleavage of the factor Xa recognition site should increase this distance greatly, leading to a marked decrease in FRET efficiency. Following assembly and HPLC purification, we suspended the substrate in a Tris-based cleavage buffer at a final concentration of 100 nM and incubated it with varying concentrations of factor Xa for 2 h. We then excited the samples using Cy3 excitation filters (540/25nm) and measured Cy3 (590/35 nm) and ATTO 643 (680/30 nm) emission via plate reader (**Methods**). We characterized the rate of substrate cleavage in terms of the FRET ratio, which is the ratio of acceptor fluorescence relative to the sum of fluorescence from both the donor and acceptor, or (*F*_*acceptor*_)/(*F*_*acceptor*_ + *F*_*donor*_). We normalized this metric by dividing the FRET ratio for a given time point from three replicate experiments by the average background FRET ratio obtained with factor Xa-free control samples at the same time point. This normalization process corrects for any baseline changes in fluorophore emission over time (**SI, Fig. S2**). Based on the Michaelis-Menten equation and associated V_max_ definition, we expect that the initial reaction rate (V_0_) should be linearly proportional to enzyme concentration given that we used a fixed substrate concentration. By performing a linear regression, we were able to confirm this and validate that our substrate can be used for the quantitative analysis of factor Xa activity (**Fig. 2**).

**Figure 2.**
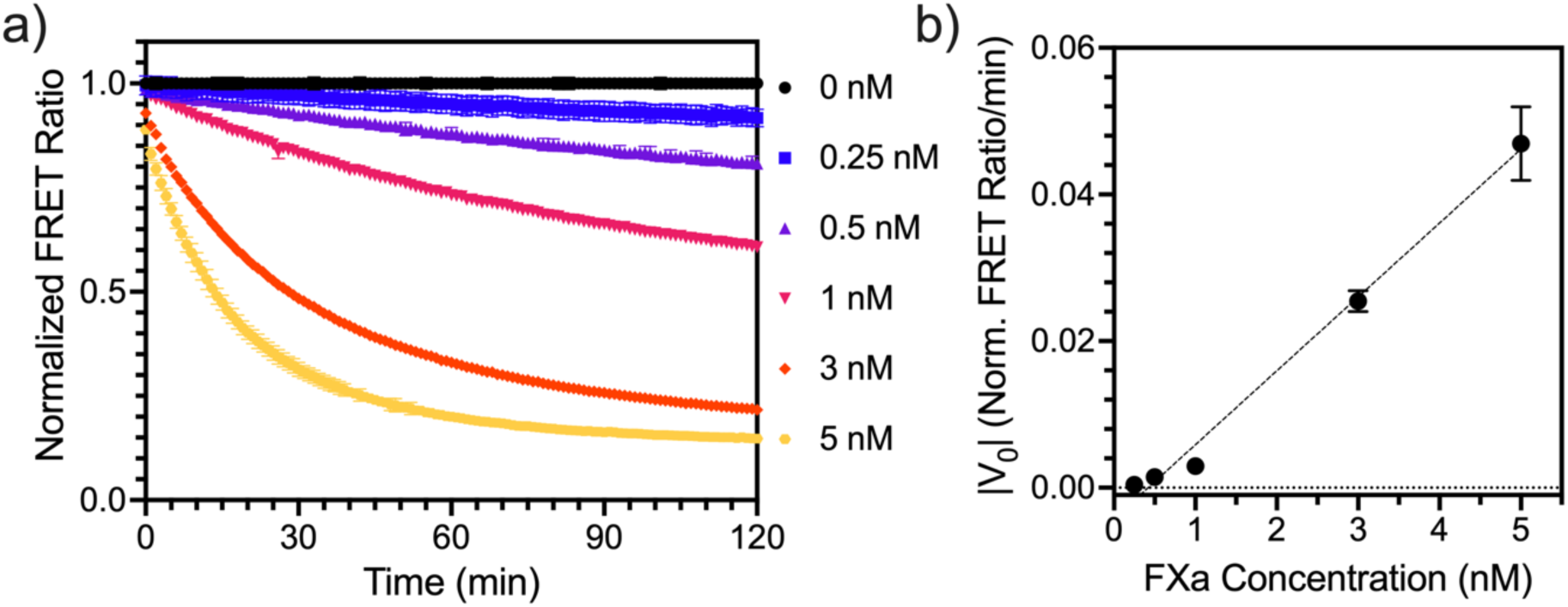
Assessing the quantitative performance of our FRET-labeled factor Xa substrate. **a**) We incubated 100 nM substrate with known concentrations of factor Xa (0 – 5 nM) in Tris-based buffer. Measurements were performed in triplicate and normalized relative to the background signal from the enzyme-free control. **b**) The initial slope (|V_0_|) of the FRET ratio curves exhibits linearity relative to factor Xa concentration (slope = 0.010 normalized FRET ratio/min/nM, R^2^ = 0.98). Plot shows mean |V_0_| values from triplicate experiments, and error bars represent the standard deviation. Mean |V_0_| and standard deviation for all concentrations are reported in **SI, Table S2**.

### On-fiber factor Xa detection

We next incorporated the FRET substrate into our evanescent sensing system to see whether it can achieve quantitative performance in a protein-rich artificial serum medium that has ionic strength, albumin content, and gamma globulin content that are designed to match those of human plasma [28, 29]. Fibers surfaces are prepared as previously described [25] and then incubated with 50nM factor Xa substrate in buffer (**Methods**). We prepared stocks of factor Xa at concentrations of 100 nM, 50 nM, 10 nM, 5 nM, and 1 nM in artificial serum. The prepared fibers were incubated directly in these various factor Xa-spiked artificial serum samples at 37 °C for 2 h while we monitored the resulting FRET ratio changes in real time (**Fig. 3a**). To facilitate comparison across sensors, donor and acceptor signals are normalized to their respective initial values prior to calculating the FRET ratio, and the resultant ‘fractional FRET ratio’ (FFR) is plotted. For comparison, we also collected measurements from a standard chromogenic assay for factor Xa activity, which we performed on the same samples (**Fig. 3b**). We measured each known concentration in triplicate using both methods and then correlated the initial slope values for each sample (**Fig. 3c**). Notably, the V_0_ values obtained from fiber-based measurements correlated closely with factor Xa concentration (r = 0.9940, R^2^=0.9881, p < 0.0001), with reasonable reproducibility across samples of the same concentration (**SI, Table S3**). The V_0_ values obtained from fiber-based measurements also correlated well with V_0_ values obtained from the chromogenic assay (r = 0.8428, R^2^=0.7102, p < 0.0001). Notably, the chromogenic assay procedure required an 80-fold dilution of artificial serum samples into a Tris-based buffer to minimize optical artifacts, while the fiber-based measurement required no such dilution. These results show that our sensor can deliver comparable quantitative performance to existing methods in protein-rich samples without the additional processing required by existing methods.

**Figure 3.**
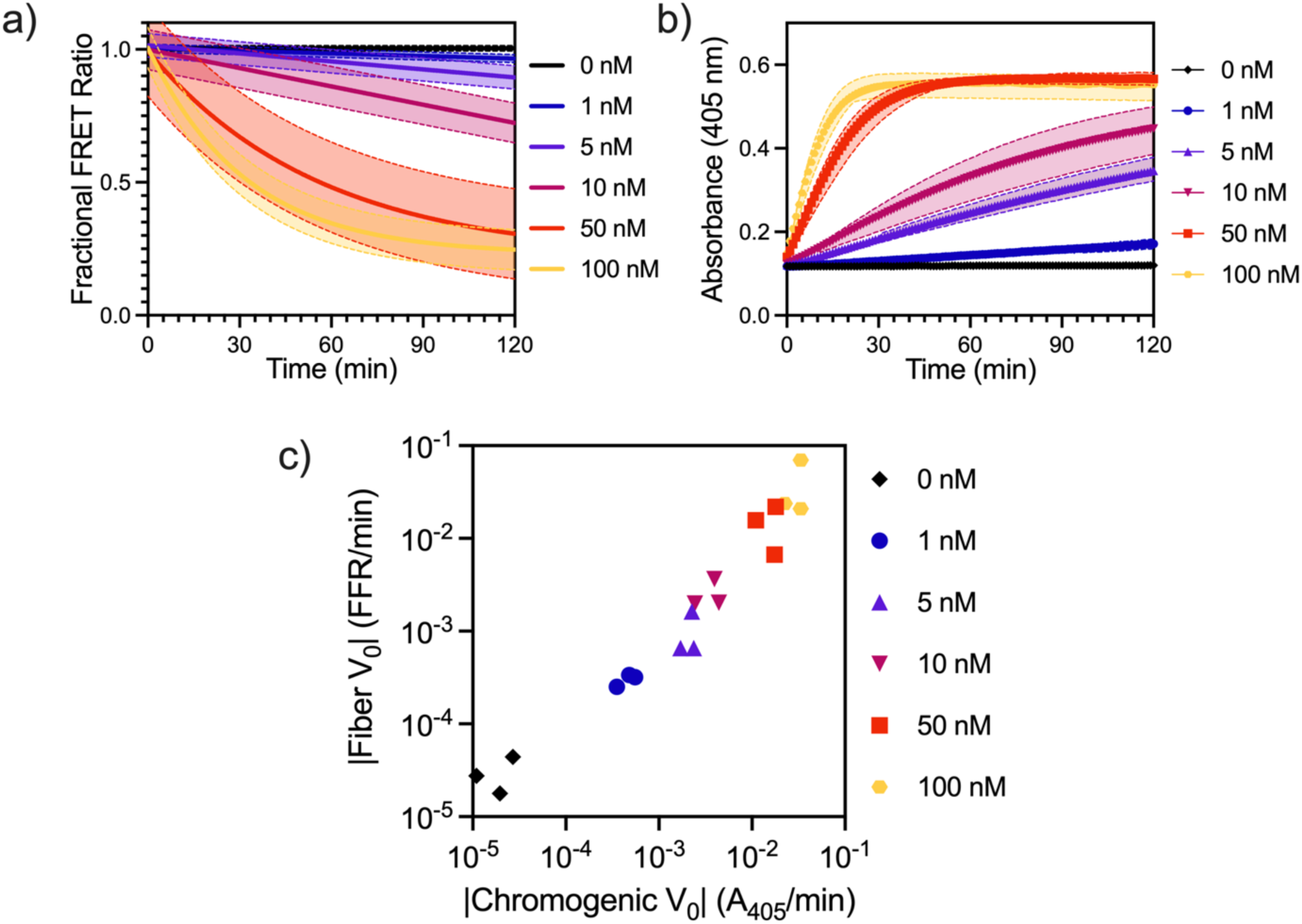
Assessing the performance of our FRET substrate on a fiber optic probe in a protein-rich environment. **a)** Detection of known factor Xa concentrations in artificial serum with fiber optic probe. Each concentration measurement was repeated three times. Data was fit to an exponential decay treating each replicate independently, and the nonlinear fit of all three replicates is presented with 90% prediction intervals (**Methods**). Due to the real-time readout, each replicate contains data from unique timepoints, making it impossible to plot conventional error bars. Raw data from triplicate experiments are presented in **SI Fig. S3**. **b)** Measurements performed with the same samples via gold-standard chromogenic assays on a plate reader. Mean values are plotted, and error bars represent the range of all replicates. Raw data from triplicate experiments are presented in **SI Fig. S3**. **c)** Correlation of V_0_ obtained via chromogenic assay (x-axis) and fiber-based assay (y-axis) (r = 0.8428). Individual results are shown from triplicate experiments

### Factor Xa detection in minimally processed human blood

We next demonstrated that our system can directly assess factor Xa activity in minimally processed human blood. Blood was collected from a single donor into a blue top sodium citrate tube and split into two aliquots. In the first, ‘inhibited’ sample, factor Xa activity was fully inhibited by spiking in sodium citrate to a final added concentration of 3% w/v (116.3 mM) prior to measurement. Citrate chelates calcium, a necessary cofactor for factor Xa activity, and 3% citrate was sufficient to effectively inactivate factor Xa for this sample. In the second, ‘non-inhibited’ sample, we instead added an equivalent volume of water to match the dilution from the added citrate. Immediately prior to measurement, calcium was added to both samples to a final concentration of 8 mM. In the ‘inhibited sample’, this additional calcium was chelated by the added citrate and unable to function as a cofactor for factor Xa. In the ‘non-inhibited sample’, the added calcium was sufficient to overcome the citrate added during blood collection and promote coagulation by functioning as a cofactor for factor Xa. We then incubated our substrate-coated fiber probes in 50 µL of each blood sample (final concentration ∼85% whole blood by volume) at 37 °C for 1 h, with measurements performed in triplicate. Within the first half hour of measurement, the FFR in the non-inhibited samples dropped to ∼0.2, indicating rapid cleavage of the substrate. In contrast, the FFR in the inhibited samples remained ∼1, indicating minimal cleavage of the substrate (**Fig. 4a**). These results verify that our system can detect endogenous factor Xa activity in minimally processed human blood within a short time frame (<30 min).

**Figure 4.**
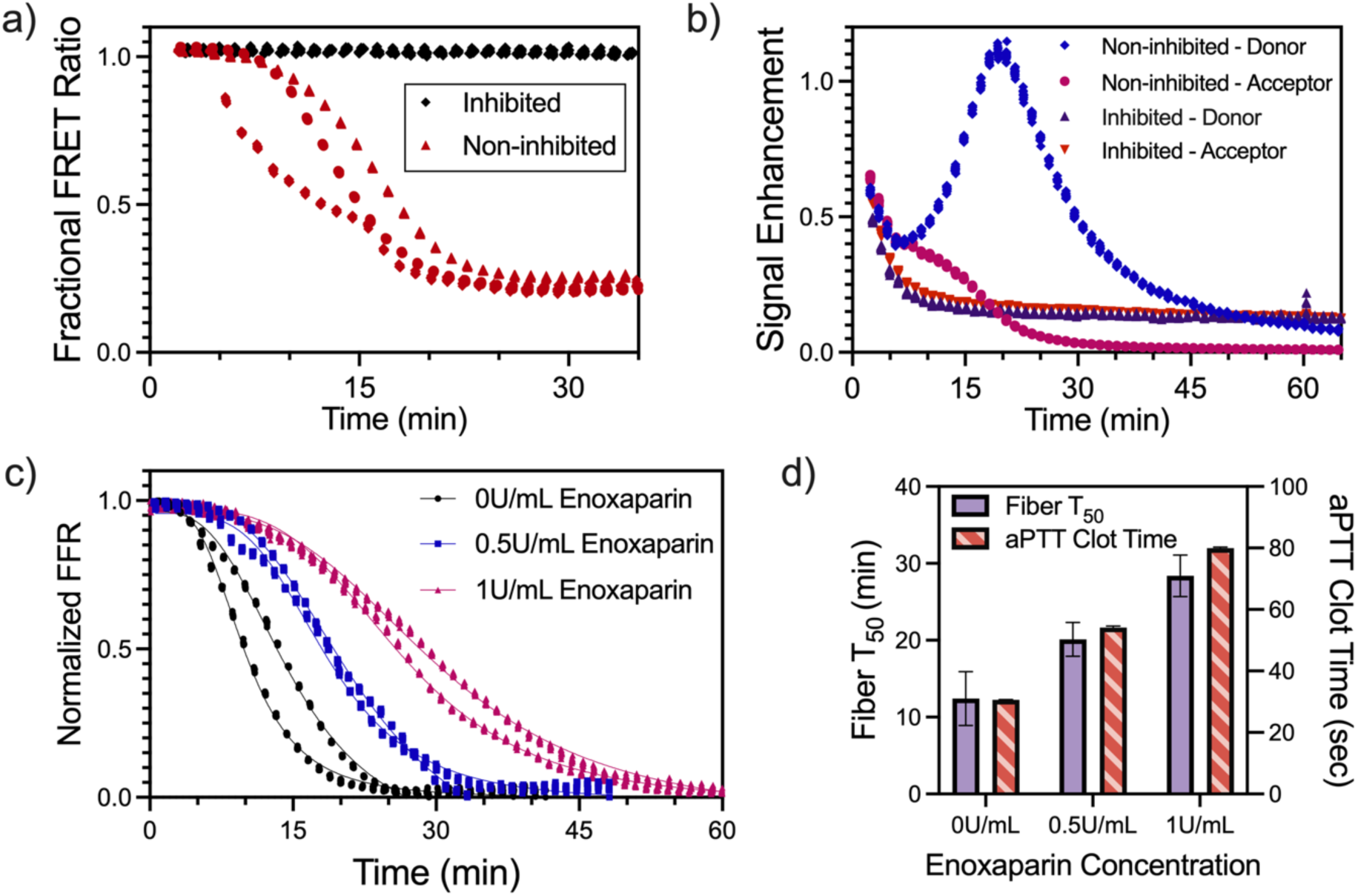
Assessing sensor performance directly in whole blood samples. **a)** Non-inhibited (red) and inhibited (black) samples of factor Xa activity in human blood. The inhibited samples show no decrease in fractional FRET ratio over time indicating no factor Xa activity. In contrast, the non-inhibited samples show a decrease in fractional FRET ratio indicating factor Xa activity. **b)** Representative data from donor and acceptor channels are shown for a single replicate of non-inhibited and inhibited samples. Notably, the inhibited sample experienced a significant signal decrease in both channels while the FRET ratio remained stable. The non-inhibited sample exhibited a significant increase in the donor channel, indicating a recovery of fluorescence emitted rather than energy transfer via FRET. **c)** Known concentrations of enoxaparin are added to a single donor sample, and normalized FFR response is plotted for duplicate measurements. For each replicate, data is plotted from the start of measurement until the minimum normalized FFR was reached to allow visual comparison of T_50_ (**SI Note 2**). **d)** T_50,_ the time to 50% of normalized FFR, is correlated to aPTT clot time, with excellent correlation (r = 1.00). Measured T50 and aPTT clot times are reported in **SI Table S4**.

Ratiometric FRET measurement is a critical feature of our system that eliminates environmental effects which confound single fluorophore measurements in complex media. In the inhibited samples, both the donor and acceptor channels exhibited notable changes in signal due to changes in the local environment of the fluorophores when blood was added to the sensor chambers (**Fig. 4b**). However, these changes were proportional across both channels, resulting in a stable FFR. In a single-color measurement, these changes in fluorescence would not be discernable from true cleavage events. In the non-inhibited samples, we observed a clear increase in donor signal over the first 20 minutes, indicating that cleavage of the substrate prevented FRET and instead resulted in donor-only fluorescence. Together, these results confirm that the combined use of evanescent detection and ratiometric FRET-based measurements can overcome the potential confounding effects of interferents in blood, allowing accurate detection of factor Xa activity.

Finally, we demonstrated that we could detect the effects of therapeutic concentrations of enoxaparin on factor Xa activity directly in minimally processed whole blood with our sensor. A citrated sample of blood from a single donor was split into three aliquots, to which we added enoxaparin at final concentrations of 0, 0.5, or 1 U/mL. The 0.5 and 1 U/mL concentrations respectively represent the lower and upper bounds of the recommended therapeutic range for both adult and pediatric patients [30, 31, 32]. We added calcium to each aliquot (final concentration = 8 mM) to overcome the sodium citrate added during blood collection and promote coagulation activity, such that the final concentration of whole blood was ∼90% by volume, and incubated 50 µL of each sample in duplicate with prepared fiber probes for 2 h at 37 °C. FFRs were calculated as described above. Due to differences in the magnitude of signal change across fibers, FFRs were normalized prior to plotting to facilitate visual comparison (**SI, Fig. S4**). We fitted these normalized FFR responses to a sigmoid and calculated the time to achieve 50% maximum signal change (T_50_) (**Fig 4c, SI Note 2, Methods**). T_50_ increased linearly with the enoxaparin concentration (r = 0.9998, R^2^=0.9996, p = 0.0129), confirming that our sensor can differentiate the inhibition of factor Xa from varying therapeutic concentrations of enoxaparin. For comparison, we performed aPTT tests with plasma samples derived from the same 90% blood samples measured using our fiber-optic sensor to gauge the downstream anticoagulant effects of enoxaparin and found similar linearity between clotting time and enoxaparin concentration (r=0.99, R^2^ = 0.99, p = 0.0175). Additionally, T_50_ values from our sensor and aPTT-derived clotting times demonstrated excellent correlation (r = 1.00, R^2^ > 0.99, p = 0.0046) **(Fig. 4d, SI Table S4, Methods**). We repeated the fiber optic and aPTT measurements with a blood sample from a second donor with enoxaparin concentrations of 0 and 0.5 U/mL (**SI, Fig. 5**). We observed similar trends between T_50_, enoxaparin concentrations, and aPTT times but with higher variance in the 0.5 U/mL enoxaparin T_50_ replicates. This variance highlights the need to use replicates for increased measurement confidence, though the consistent trend across donors is a promising indication of our system’s sensitivity to anticoagulants. Together, our results demonstrate the ability to measure physiologically relevant, specific molecular change at the point of drug action in minimally processed blood that directly propagates to downstream clotting outcomes measured with traditional clotting assays in plasma.

## Conclusions

In this work, we have demonstrated the ability to directly measure factor Xa activity in minimally processed whole blood and shown that this detection is comparable with the results of existing gold-standard clinical assays that require prior processing of blood into plasma. Our sensor design, which exploits an evanescent field for fluorescence detection and a ratiometric FRET-based readout, enables us to overcome the confounding effects of both high protein content and red blood cells. We have demonstrated that we can sensitively discriminate the response to various therapeutic concentrations of enoxaparin, a LMWH routinely used in pediatric patients, in small sample volumes (50 µL per replicate) of minimally processed blood. Critically, we have consistently shown that the measurements of factor Xa activity produced by our sensor correlate closely with both chromogenic assays for factor Xa activity and measurements of downstream coagulation time in the aPTT assay.

Our sensor could prove useful for performing point-of-care assessment and monitoring of coagulation therapies with the numerous antithrombotic drugs that target factor Xa activity. By measuring at the site of drug action, our sensor provides specific molecular information for a new window of insight, enabling better guidance for tasks like enoxaparin dosing in pediatric populations utilizing reduced whole blood volumes. Additionally, given factor Xa’s role in the common pathway of the coagulation cascade, our sensor could also be used a general tool to measure overall coagulation function in point-of-care settings with minimal sample preparation. Further studies and characterization will be needed before the sensor is appropriate for use in a clinical workflow. Establishing a workflow that minimizes variance between replicate measurements is necessary to overcome the variance observed in the replicate measurements taken from the additional donor sample. Our sensor will need to be tested across larger cohorts of donors to validate its sensitivity, identify potential sources of variance and error seen in cross-donor replicates, and establish the therapeutic range for T_50_ values as is done for aPTT assays to account for baseline individual variance. Clinical workflow, wherein blood samples would not necessarily be spiked with anticoagulant upon collection but rather injected directly into the sensor, would also need to be established and validated. These issues will be the focus of future research, but we believe the results presented indicate new opportunities to leverage the direct detection of factor Xa activity in whole blood to guide the dosage of anticoagulant drugs to mitigate bleeding and other complications associated with anticoagulant therapy.

## Materials and Methods

### Reagents and materials

All chemicals, buffers, and buffer components were ordered from Fisher Scientific unless otherwise specified. The DNA substrate sequence was ordered from IDT with HPLC purification, and the substrate peptide was ordered from GenScript (see **SI, Table S1**). Cy3 N-hydroxysuccinimide (NHS) ester was ordered from Lumiprobe (product no. 11020). Atto643 NHS Ester was ordered from Atto-Tec (product no. AD 643). Factor Xa was ordered from New England Biolabs (product no. P8010L). Plate-reader half-area black flat-bottom polystyrene 96-well plates were ordered from Corning (product no. 3694) for custom factor Xa substrate experiments and clear, flat bottom polystyrene 96-well microplates from Corning (product no. 9017) were used for chromogenic assays. Cleavage buffer was prepared with 20 mM Tris, 1 mM CaCl_2_, and 50 mM NaCl at pH 7.5. Human serum albumin (HSA) was purchased from VWR International (product no. 10148-964). Human serum gamma-globulin (HSG) was purchased from Sigma Aldrich (product no. 345886). Artificial serum was adapted from references [28, 29] and prepared with 142 mM NaCl, 2.5 mM CaCl_2_, 1.5 mM MgCl_2_, 5 mM KCl, 4.2 mM NaHCO_3_, 50 mM Tris, 42 mg/mL HSA, and 1.6 mg/mL HSG with a pH of 7.4. The factor Xa chromogenic substrate was purchased from Sigma (product no. F3301). Sodium citrate was ordered from Fisher Scientific (catalog no. 18-606-449). Citrated healthy human donor whole blood was obtained from the Stanford Blood Center on the same day that fiber-optic measurements were performed. Blood was stored at RT until 30 min prior to measurement. Enoxaparin sodium was purchased from Sigma-Aldrich (product no. 1235820). Factor Assay Control Plasma (product no. 0020-1) and factor X-deficient plasma (product no. 1000) were obtained from George King Bio-Medical, Inc.

### Instrumentation

HPLC purification was carried out on an Agilent 1260 Infinity II using a PRP-1 reversed-phase HPLC column (Hamilton, product no. 79425). HPLC utilized acetonitrile (ACN) from Fisher (product no. A996-4) as an organic solvent and 50 mM triethylammonium acetate (TEAA) from Glen Research (product no. 60-4110-52) as an ion pairing reagent. The method used was as follows: two minutes at 95% 50mM TEAA and 5% ACN followed by a ramp at constant gradient over 28 minutes to final proportions of 50% 50mM TEAA and 50% ACN. Spectrophotometry measurements were taken on a NanoDrop 2000 (Thermo Fisher Scientific). Plate-reader measurements were taken on a Synergy H1 hybrid microplate reader (Agilent BioTek). FRET measurements on the plate reader were taken with a Cy3-Cy5 two-channel filter cube (Agilent BioTek, Cat. No. 1035106). The aPTT assay was performed using a Start4 coagulation analyzer (Stago International, Asnières-sur-Seine, France).

### Factor Xa substrate synthesis

Substrate DNA strands were prepared by labelling the internal amine group of the substrate DNA strand with Cy3 NHS ester. Labelling was carried out by mixing 240 µL of 2 µM DNA with 30 µL of sodium bicarbonate solution (7.5% w/v), 1.5 µL of 50 mM Cy3-NHS, and 7.5 µL dimethylformamide (DMF) for ester solubility, and then incubating overnight at room temperature. Free Cy3-NHS was removed via ethanol precipitation, followed by resuspension in 25 µL deionized water. The resuspended DNA was then HPLC-purified to remove unlabeled DNA, lyophilized and then resuspended in DI water and stored at −20 °C prior to peptide conjugation. The substrate peptide was first resuspended in anhydrous DMF at 25 mM. Labeling was carried out by mixing 30 µL of 20 mM Atto643 NHS in DMF, 16 µL of 25 mM resuspended peptide, and 1.2 µL of N,N-diisopropylethyamine and incubating overnight at room temperature. To assemble the full DNA-peptide conjugate substrate via click chemistry, 7 µL of Atto643-labeled peptide (which contained a C-terminal azide group) was mixed with 28.5 µL of 119 µM Cy3-labeled DNA (which contained a 5’ dibenzocyclooctyne group) and incubated at room temperature overnight. The reaction was then HPLC purified, and the full substrate construct was purified by identifying the elution peak with the expected UV absorption at A260/A280 (DNA), A569 (Cy3) and A643 (Atto643) values (**SI, Fig. S6**). After elution, the full substrate was lyophilized, resuspended in deionized water to a final concentration of 10 µM, and stored at 4 °C.

### Plate-reader validation of the factor Xa substrate

2 µL of 2 µM substrate in cleavage buffer was pipetted into 34 µL of cleavage buffer in a 96-well plate, with triplicate experiments for each tested concentration of factor Xa (0, 0.25 nM, 0.5 nM, 1 nM, 3 nM, 5 nM). The plate was then incubated in the plate-reader at 37 °C for 10 min to thermally equilibrate. During plate equilibration, factor Xa was serially diluted in cleavage buffer to obtain stocks of 50 nM, 30 nM, 10 nM, 5 nM, and 2.5 nM. After equilibration, 4 µL of the appropriate Xa stock was immediately added to and pipette-mixed with the substrate-containing wells (4 µL of cleavage buffer was used for 0 nM Xa). The plate was then analyzed at 37 °C on the plate-reader for 2 h, with FRET measurements taken every minute. The resultant data were normalized to the average value of the 0 nM case. The dynamic range of this assay was calculated by fitting the 5 nM factor Xa data to an exponential decay (Y = (Y0 - Plateau)*exp(-K*X) + Plateau) and subtracting the plateau value from the starting value of 1. The initial rate (V_0_) was calculated as the slope over the time taken for the magnitude of signal change to reach 10% of the dynamic range, with a minimum of 3 time points used.

### Optical Readout Hardware

Tapered fibers were produced as described in reference [25]. The optoelectronic readout hardware was adapted from reference [25] with modifications to allow two color measurement and time multiplexed measurements of multiple tapered fiber probes. A Coherent Obis 532-20 LS (532 nm laser, 20 mW) was used for fluorescence excitation and driven by an Obis LX/LS Scientific Remote (Part Number: 1234466). The laser was input to a Multimode MEMS fiber to fiber 1×2 switch (Agiltron FFSW-123872332, GIF 62.5/125, FC/PC connectors) that was controlled by a switch driver (Agiltron SWDR-111111121). This switch allowed for alternate use of the 532 nm laser or another excitation source, though 532 nm excitation was exclusively used in this work. The 532 nm source was filtered using a neutral density filter (Chroma ND 1.0 - 10% Trans V2) and 532 nm bandpass filter (Thorlabs FL532-3) prior to being input to a Multimode MEMS fiber to fiber 1 x 8 switch (Agiltron FFSW-183572332, GIF 62.5/125, FC/PC connectors) that was controlled by a switch driver (Agiltron SWDR-58A241121). 7 output ports of this switch were attached to GIF 625 fiber pigtails (Thorlabs) and available for splicing in tapered fiber probes. The final output port was used to monitor laser power fluctuations over time via a Thorlabs PM100D power meter. During all experiments, each port was sequentially selected, such that the FRET signal from each probe could be measured. After all samples were measured, the final port was selected to measure power fluctuations over time, and the process was repeated. Collected signals from each probe were returned through the switch and along an emission pathway. This pathway was split using a 1×2 fiber splitter (Thorlabs TG625R2F1A) into a donor and acceptor pathway. The donor emission pathway was filtered using a 550 nm long pass filter (Thorlabs FELO550) and 595 nm band pass filter (Chroma ET595/50m), and the acceptor emission pathway was filtered using a 676 nm band pass filter (Semrock BrightLine FF01-676/29-25). Both channels were then measured using SPCMs (Excelitas SPCM-AQRH). For the final port used to monitor power, light reflected from the power meter photodiode was measured by the SPCMs. In this work, the excitation power at the output ports of the 1×8 switch was approximately 500nW. Measurements were taken with an integration time of 500ms per data point, and 15 data points were collected per fiber probe before switching to the next fiber probe. Data was collected using a custom LabVIEW GUI.

### Fiber Data Processing

After data collection, data was scaled to account for power fluctuations over time measured using the recorded SPCM counts from the final 1×8 switch port. Prior to substrate functionalization, a background scattering signal was measured for each fiber probe in cleavage buffer. The average background signal for donor and acceptor channels was calculated by averaging over at least 75 data points and subtracted from all collected data on the respective channels. After substrate functionalization, a baseline signal was measured in cleavage buffer for each fiber probe, with at least 120 points collected per fiber. The average baseline signal was calculated per channel, and the donor and acceptor channels were normalized to their respective baselines to calculate signal enhancement. The donor and acceptor signal enhancements were subsequently used to calculate FFR. After baseline was collected, samples of interest were added to each probe for measurement.

### Validation of factor Xa substrate on our fiber-optic sensor in artificial serum

Fiber-optic probes were constructed and prepared as detailed in previous work [25], up until the injection of neutravidin after the fibers were sealed in chambers. Chambers containing fibers were filled with 40 µL of 0.2 mg/mL neutravidin in cleavage buffer and incubated for 10 minutes at room temperature. Chambers were then washed twice with 200 µL of cleavage buffer to remove unbound neutravidin. Measurements were collected at this point for background subtraction. Factor Xa substrate was prepared as specified above, and 40 µL of 50 nM substrate in cleavage buffer was injected into the fiber chambers and incubated for 30 minutes at room temperature. The chambers were then washed twice with cleavage buffer to remove unbound substrate and placed in an incubator at 37 °C. After five minutes, sensor signal was measured to determine baseline. Factor Xa was serially diluted to make stocks at 100 nM, 50 nM, 10 nM, 5 nM, and 1 nM in artificial serum. Fibers in chambers and prepared factor Xa stocks were both incubated at 37 °C for 10 minutes prior to measurement. Following this incubation, 50 µL of factor Xa stocks were injected into fiber chambers, and fibers were measured approximately every minute for an hour to observe factor Xa activity. The recorded FRET signals were background subtracted, normalized, and used to compute fractional FRET ratio change over time as described above. For plotting purposes, the 0 time point was assigned as the midpoint in time between the final baseline measurement point and the first measurement point in the artificial serum samples. The dynamic range of this assay was calculated by fitting the 100 nM factor Xa data to an exponential decay (Y = (Y0 - Plateau)*exp(-K*X) + Plateau) and subtracting the plateau value from the starting value of 1. The initial rate (V_0_) was calculated as the slope over the time taken for the magnitude of signal change to reach 10% of the dynamic range.

Concurrently, we performed a chromogenic assay on the same factor Xa stocks in triplicate, and measured absorbance at 405 nm using a plate reader. Briefly, 2 µL of 20 mM chromogenic substrate in cleavage buffer was added to 77 µL of cleavage buffer in wells and incubated at 37 °C in the plate reader for 10 min. Then 1 µL of factor Xa stocks were added to the wells to achieve an 80X dilution per the manufacturer’s recommendation. Absorbance at 405 nm was measured every minute at 37 °C for 2 h. The dynamic range of this assay was calculated by fitting the 100 nM factor Xa data to a one phase association (Y = Y0 + (Plateau-Y0)*(1-exp(-K*x))) to find the plateau value. The initial rate (V_0_) was calculated as the slope over the time taken for the magnitude of signal change to reach 10% of the plateau value.

### Factor Xa activity detection in minimally-processed whole blood

Optical fiber probes were prepared as above, including incubation with the factor Xa substrate and subsequent washing. The fiber probe chambers were then injected with 50 µL artificial serum and incubated for 1 h at RT to passivate the chamber surface before washing with 200 µL cleavage buffer and storage at room temperature. Citrated healthy human blood samples were collected on the day of measurement. Two 900 µL aliquots were pipetted from each blood sample. The ‘inhibited’ aliquot was mixed with 100 µL of 30% (w/v) sodium citrate, and the ‘non-inhibited’ aliquot was mixed with 100 µL deionized water. Both samples were then briefly vortexed and incubated at room temperature for 30 min on a rotor. 285µL of each sample was removed and spiked with 15 µL of 160 mM CaCl_2_ to reverse the effect of citrate spiked in during blood collection. 50 µL of each recalcified fraction was injected into fiber chambers, and fiber chambers were measured at 37 °C for 2 h to monitor factor Xa activity. The recorded fluorescence signals were background subtracted, baseline normalized, and used to compute fractional FRET ratio change over time as described above. The 0 time point was assigned at 2 minutes prior to the first measurement point in blood samples to account for the time required to load samples into chambers, place chambers in the incubator, and begin the measurement.

### Factor Xa detection in minimally-processed whole blood treated with enoxaparin

Optical fiber probes were prepared as specified in the prior section. Healthy human blood samples were collected on the day of measurement. Enoxaparin was serially diluted in deionized water to create 20 U/mL and 10 U/mL stocks. Aliquots of 855 µL from each blood sample were mixed with 45 µL of the 20 U/mL enoxaparin stock, 45 µL of the 10 U/mL enoxaparin stock, and 45 µL of deionized water to create blood samples containing 1.0 U/mL, 0.5 U/mL, and 0 U/mL enoxaparin, respectively. All aliquots were briefly vortexed and incubated at room temperature for 30 min on a rotor. 190 µL of each sample was removed for fiber-based measurement. Each 190 µL fraction was spiked with 10 µL of 160 mM CaCl_2_ immediately prior to injecting 50 µL of each aliquot into the fiber chambers. The fiber chambers were incubated at 37 °C for 2 h to measure factor Xa activity. The recorded fluorescence signals were background subtracted, baseline normalized, and used to compute fractional FRET ratio change over time as described above. As before, the 0 time point was assigned at 2 minutes prior to the first measurement point in blood samples to account for the time required to load samples into chambers, place chambers in the incubator, and begin the measurement. Data from time 0 until the minimum value data point was extracted and used for subsequent analysis and plotted in **Figure 4c**. This FFR data was normalized and fit to a sigmoid curve, and the time to achieve 50% maximum signal change (T_50_) was calculated.

The remaining volume of samples (not yet recalcified) were used to perform the one-stage clotting-activated partial thromboplastin time (aPTT) assay in duplicate to assess normal coagulation activity of the intrinsic coagulation pathway factors through a viscosity-based detection system. Briefly, blood was spun down at 3500 RPM for 15 minutes, aliquoted, and stored at −80°C. Plasma was thawed and warmed to 37°C for 8 minutes prior to assessing clot times in duplicate on a Start4 coagulation analyzer according to the manufacturer’s instructions for an aPTT assay. Factor Assay Control Plasma containing 118% factor X activity and factor X-deficient plasma provided a normal and abnormal control, respectively.

Correlation of T_50_ and aPTT values was performed in GraphPad Prism using duplicate measurements presented in **SI Table S4**. Using this correlation function, the mean values of duplicate measurements were found and subsequently used to calculate correlation.

## Acknowledgements

This work was supported by The Helmley Trust and Wellcome LEAP SAVE program. A.P.C. acknowledges support from the NSF Graduate Research Fellowship Program and the Stanford Graduate Fellowship. B.C.W. acknowledges funding from the Knight-Hennessy Scholars Program and the NSF Graduate Research Fellowship. S.Y. acknowledges support from the Stanford Graduate Fellowship. H.C.C. is funded by the National Bleeding Disorders Foundation Judith Graham Pool Postdoctoral Research Fellowship. G.B. is funded by NHLBI grant K99HL150595, Hemophilia of Georgia Clinical Scientist Development Award, Duda Foundation, and is a Anne T. and Robert M. Bass Endowed Faculty Scholar in Pediatric Cancer and Blood Diseases. The authors would like to thank Dr. Amani Hariri, Dr. Brian Young, Dr. Yasser Gidi, Dr. Tuan Trinh, and Grace Maddocks for their insightful discussion and assistance and all members of the Soh lab for their support throughout this project. We would also like to thank Professor Michel Digonnet’s lab for their technical assistance in producing tapered fiber probes.

