## Supplementary Information for "Direct Optical Detection of Factor Xa Activity in Minimally Processed Whole Blood"

**SI Note 1**

Peptide design is adapted from [33, 34]. Given the significant fluorescence change seen in [33], we anticipate the scale of our substrate to be appropriate for significant FRET ratio change upon cleavage. Our chosen sequence was guided by characterization performed in [34], with additional focus on introducing hydrophilic residues unlikely to interfere with other residues in the sequence.

**SI Note 2**

Due to feedback activation within the coagulation cascade, we expect the FRET ratio change in non-inhibited blood samples, including samples partially inhibited by therapeutic concentrations of enoxaparin, to be sigmoidal. This behavior precludes the use of $V_0$ as a metric. For this reason, we use $T_{50}$ to characterize the timescale of coagulation, providing an indicator for the degree of inhibition accomplished by enoxaparin. After substrate cleavage is largely completed, we hypothesize that additional factors such as photobleaching and additional environmental effects begin to compete with residual substrate cleavage, resulting in asymmetric changes in the donor and acceptor channels and a measured increase in FRET ratio (**SI Figure S4**). For clarity, data presented in Figure 4c only includes the range over which FRET ratio is decreasing.

**Table S1**

| | |
|---|---|
| ***DNA Anchor*** | /5DBCOTEG/TTT TT/iAmMC6T/ TTT TTT TTT TTT TTT TTT /3Bio/ |
| ***Peptide Substrate*** | GGSGIEGRAAYGK(N3) |

**Table S2**

| **Enzyme Concentration (nM)** | **Mean $\|V_0\|$** (Norm. FRET Ratio/min) | **St.Dev. $V_0$** (Norm. FRET Ratio/min) |
|---|---|---|
| 0.25 | 0.0004 | 0.0004 |
| 0.5 | 0.0014 | 0.00004 |
| 1 | 0.0030 | 0.0001 |
| 3 | 0.0254 | 0.0014 |
| 5 | 0.0469 | 0.0050 |

**Table S3**

| Enzyme Concentration (nM) | Mean \|Fiber $V_0$\| (FFR/min) | % CV of \|Fiber $V_0$\| | Mean \|Chromogenic $V_0$\| ($A_{405}$/min) | % CV of \|Chromogenic $V_0$\| |
|---|---|---|---|---|
| 0 | 0.00003 | 44.74 | 0.00002 | 41.83 |
| 1 | 0.00030 | 14.73 | 0.00046 | 21.93 |
| 5 | 0.00098 | 56.96 | 0.00211 | 16.66 |
| 10 | 0.00256 | 37.11 | 0.00360 | 28.60 |
| 50 | 0.01474 | 51.65 | 0.01550 | 25.19 |
| 100 | 0.03811 | 71.80 | 0.02983 | 21.29 |

**Table S4**

| Enoxaparin Concentration (U/mL) | Fiber $T_{50}$ (min) | | aPTT Clot Time (sec) | |
|---|---|---|---|---|
| 0 | 9.957 | 14.90 | 30.5 | 30.7 |
| 0.5 | 21.670 | 18.57 | 54.5 | 53.7 |
| 1 | 26.46 | 30.28 | 80.2 | 79.7 |

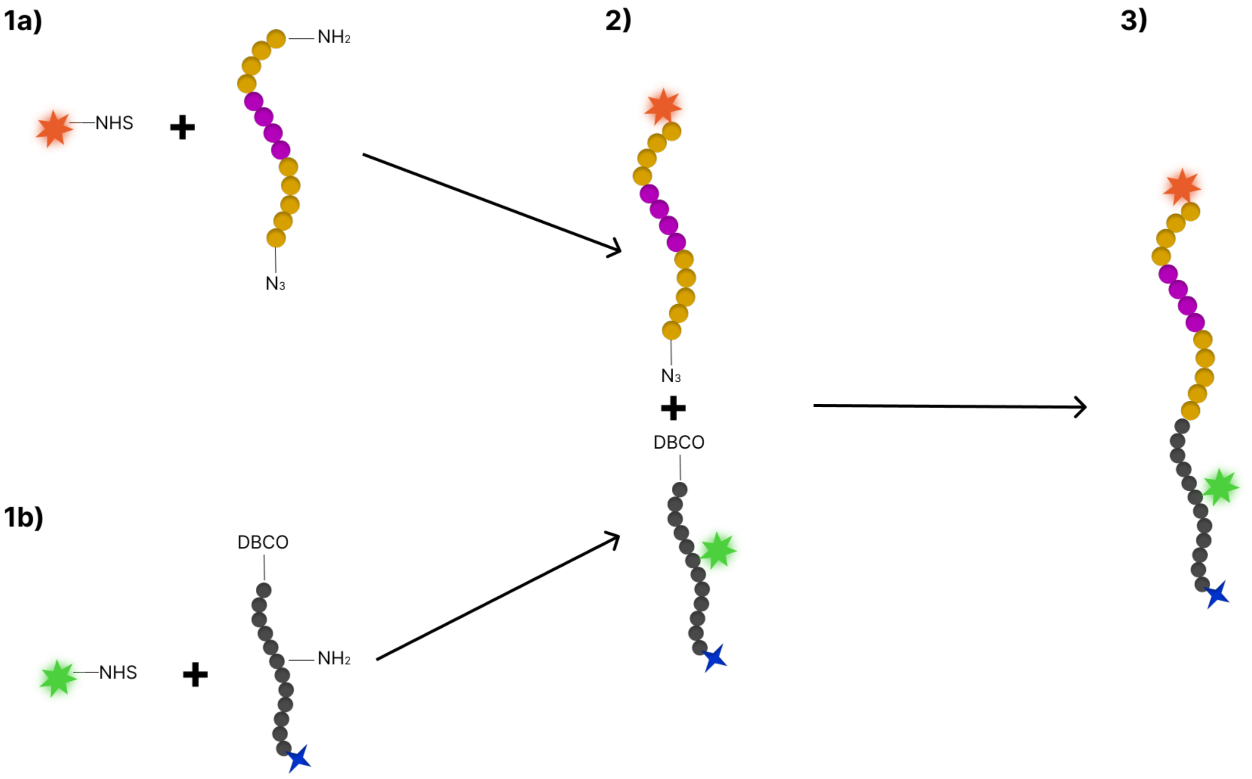


**SI Figure S1** Schematic overview of substrate synthesis process as outlined in Methods. 1a) Peptide sequence is labeled with Atto643 NHS ester at the N terminus. 1b) DNA sequence is labeled with Cy3-NHS ester at internal amine modification. 2) Labeled peptide and DNA sequences are mixed for conjugation via click chemistry between a C terminal azide and 5' DBCO. 3) Click product is full substrate with FRET pair labelling on opposite sides of factor Xa cleavage site (purple). See **Methods** for details on purification process between steps.

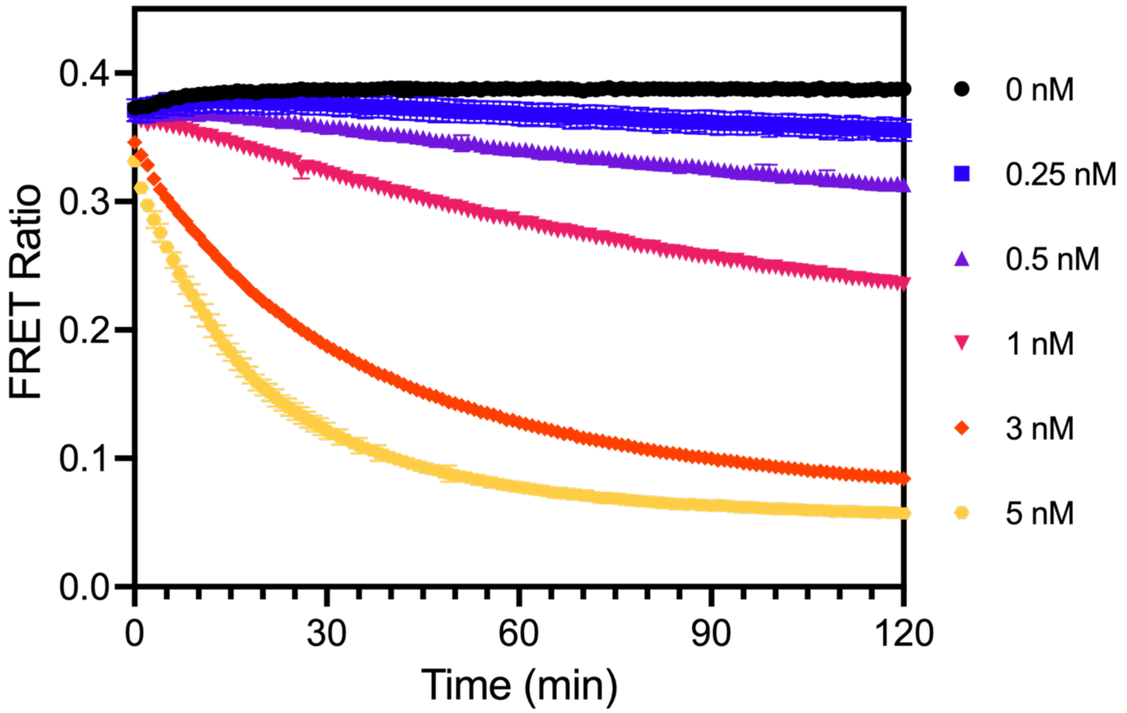


**SI Figure S2** FRET Ratio vs time prior to normalization for 0, 0.25, 0.5, 1, 3 and 5 nM factor Xa. In the 0 nM condition, Cy3 and Atto643 exhibit asymmetric changes in fluorescence over time, leading to an increase in FRET ratio. In all other conditions, the FRET ratio decreases over time. For ease of comparison, normalized data is presented in main text.

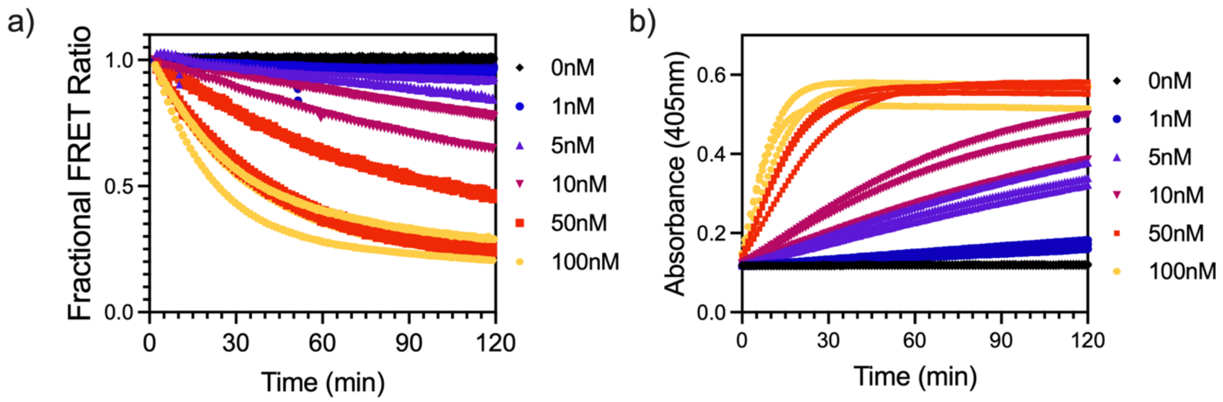


**SI Figure S3** a) Replicate data from Figure 3a. b) Replicate data from Figure 3b.

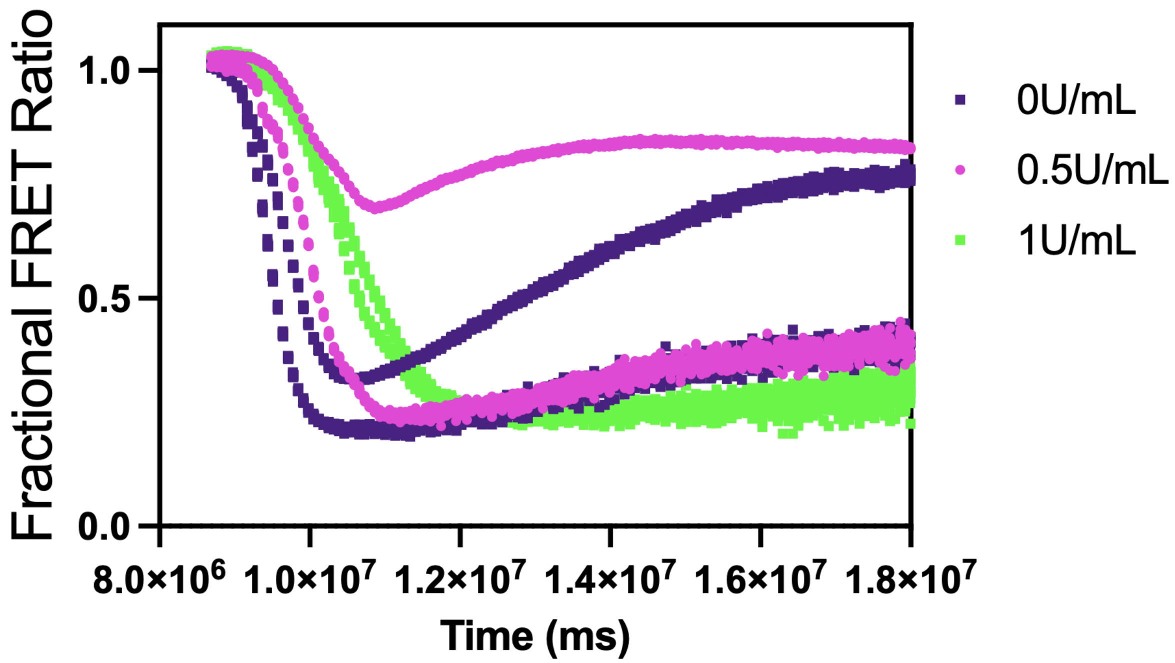


**SI Figure S4** Fractional FRET ratio for data presented in Fig 4c is plotted, with noticeable differences in the magnitude of signal change across samples. This is likely due to differences in fiber geometry or surface density of the peptide, though we consider $T_{50}$ to characterize the degree of inhibition accomplished by added enoxaparin (**SI Note 2**).

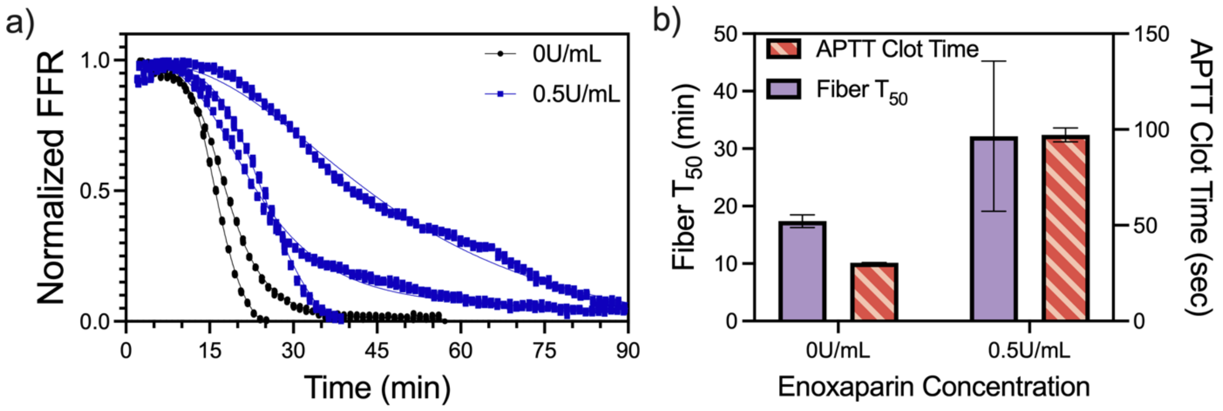


**SI Figure S5** a-b) Normalized FFR and $T_{50}$ results for an additional donor, with significant variation between the measured $T_{50}$ for 0.5U/mL replicates. Higher variance between 0.5 U/mL enoxaparin concentration replicates for fiber optic measurements leads to poorer overall correlation (r=0.66, $R^2 = 0.43$, $p = 0.23$) compared to Figure 4. Correlation between enoxaparin concentration and aPTT clotting time is similar to Figure 4 (r=0.99, $R^2 = 0.99$, $p = 0.0015$).

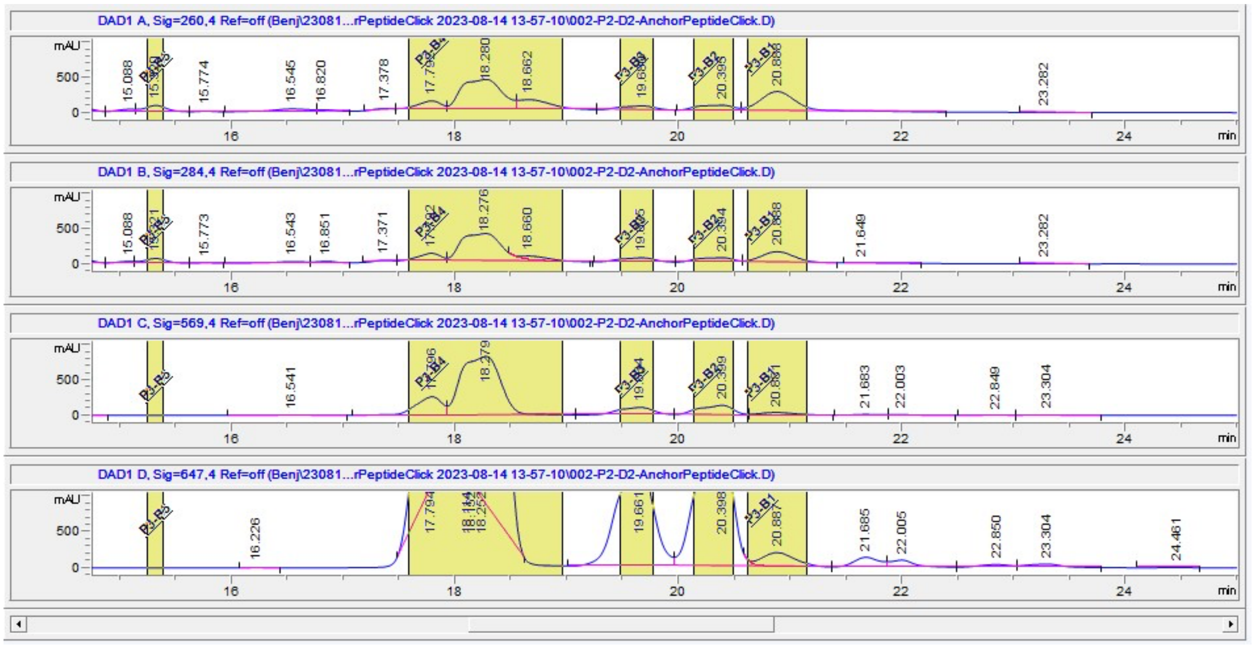


**SI Figure S6** HPLC absorption traces for purification following click conjugation of Cy3 labeled DNA and Atto643 labeled peptide. A260, A284, A569 absorption and A647 absorption traces are shown. Elution peak at 20.88 minutes was identified as the fully conjugated product. The ratio of A260 and A284 channels is as expected for the substrate (DNA absorbance is far greater than that of the peptide), A569 and A647 roughly match expected absorbance for successful Cy3 and Atto647 labelling based on dye extinction coefficients. HPLC method used described in Methods.
